# Multimodal histopathologic models stratify hormone receptor-positive early breast cancer

**DOI:** 10.1101/2024.02.23.581806

**Authors:** Kevin M. Boehm, Omar S. M. El Nahhas, Antonio Marra, Pier Selenica, Hannah Y. Wen, Britta Weigelt, Evan D. Paul, Pavol Cekan, Ramona Erber, Chiara M. L. Loeffler, Elena Guerini-Rocco, Nicola Fusco, Chiara Frascarelli, Eltjona Mane, Elisabetta Munzone, Silvia Dellapasqua, Paola Zagami, Giuseppe Curigliano, Pedram Razavi, Jorge S. Reis-Filho, Fresia Pareja, Sarat Chandarlapaty, Sohrab P. Shah, Jakob Nikolas Kather

## Abstract

For patients with hormone receptor-positive, early breast cancer without *HER2* amplification, multigene expression assays including Oncotype DX ® recurrence score (RS) have been clinically validated to identify patients who stand to derive added benefit from adjuvant cytotoxic chemotherapy. However, cost and turnaround time have limited its global adoption despite recommendation by practice guidelines. We investigated if routinely available hematoxylin and eosin (H&E)-stained pathology slides could act as a surrogate triaging data substrate by predicting RS using machine learning methods. We trained and validated a multimodal transformer model, Orpheus, using 6,203 patients across three independent cohorts, taking both H&E images and their corresponding synoptic text reports as input. We showed accurate inference of recurrence score from whole-slide images (r = 0.63 (95% C.I. 0.58 - 0.68); n = 1,029), the raw text of their corresponding reports (r = 0.58 (95% C.I. 0.51 - 0.64); n = 972), and their combination (r = 0.68 (95% C.I. 0.64 - 0.73); n = 964) as measured by Pearson’s correlation. To predict high-risk disease (RS>25), our model achieved an area under the receiver operating characteristic curve (AUROC) of 0.89 (95% C.I. 0.83 - 0.94), and area under the precision recall curve (AUPRC) of 0.64 (95% C.I. 0.60 - 0.82), compared to 0.49 (95% C.I. 0.36 - 0.64) for an existing nomogram based on clinical and pathologic features. Moreover, our model generalizes well to external international cohorts, effectively identifying recurrence risk (r = 0.61, *p* < 10^-4^, n = 452; r = 0.60, *p* < 10^-4^, n = 575) and high-risk status (AUROC = 0.80, *p* < 10^-4^, AUPRC = 0.68, *p* < 10^-4^, n = 452; AUROC = 0.83, *p* < 10^-4^, AUPRC = 0.73, *p* < 10^-4^, n = 575) from whole-slide images. Probing the biologic underpinnings of the model decisions uncovered tumor cell size heterogeneity, immune cell infiltration, a proliferative transcription program, and stromal fraction as correlates of higher-risk predictions. We conclude that at an operating point of 94.4% precision and 33.3% recall, this model could help increase global adoption and shorten lag between resection and adjuvant therapy.

## Introduction

Hormone receptor-positive disease without HER2 overexpression or amplification (HR+/HER2-) is the most common subtype of early breast cancer (EBC), accounting for approximately 70% of diagnoses^1^. A major challenge in the management of this disease has been identifying the cancers for which adjuvant chemotherapy does not meaningfully reduce the risk of recurrence. Risk stratification of HR+/HER2-EBC relies on the integration of traditional clinicopathological features (e.g., tumor size, nodal status, Nottingham grade) with multigene assays to estimate risk of recurrence and personalize adjuvant therapy. Among the commercially-available assays, the Oncotype DX (ODX) ® (Genomic Health, Redwood City, CA) is the most extensively validated and widely used in clinical practice. By measuring transcriptional abundance of 16 genes, including *ESR1*, *PGR*, *HER2*, *MKI67*, and *MMP11*, against the abundance of five reference genes using reverse transcription quantitative real-time PCR^2^, ODX calculates a recurrence score (RS) ranging from zero to 100 with both prognostic and predictive value ^2–9^.

Substantial clinical evidence from retrospective and prospective trials has shown that ODX can improve clinical decision-making in breast cancer. Retrospective analyses of the NSABP B14^2^ and TransATAC^5^ trials demonstrated the prognostic value of ODX in stratifying the risk of recurrence for HR+/HER2-EBC patients. Similarly, analyses of the NSABP B20^6^ and SWOG8814^7^ clinical trials established the predictive value of ODX by uncovering a survival benefit with the addition of adjuvant chemotherapy to endocrine therapy for patients with high risk of disease relapse. These studies provided the rationale for the prospective evaluation of ODX in the TAILORx^8^ (>10,000 patients with node-negative disease) and RxPONDER^9^ (5,083 patients with one to three positive lymph nodes) trials and established ODX as the preferred genomic assay for adjuvant treatment-decision making in HR+/HER2-EBC ^10,11^

While guidelines have recommended the use of ODX or other assays for more than a decade^10–12^, reimbursement restrictions and global accessibility barriers have limited universal adoption^13^. Beyond the United States, the cost of around 4,000 USD per sample^14,15^ and turnaround time delaying start of therapy have created barriers to adoption, despite analyses indicating downstream savings from more tailored adjuvant therapy^16^. Some efforts have been undertaken to develop nomograms based on clinical and pathologic features annotated during the standard of care, aiming to predict ODX scores ^17^. However, such tools require manual extraction of relevant inputs from the unstructured electronic healthcare record and leave room for improvement in terms of performance, with the assay itself still providing greater cost effectiveness than clinical risk tools^16^.

We investigated the use of whole-slide images (WSIs) from routinely available formalin-fixed paraffin-embedded (FFPE) tissue slides stained with hematoxylin and eosin (H&E) to predict RS. As previous studies have demonstrated, these slides can be effectively analyzed using deep learning algorithms to predict relapse risk ^18–26^. Such algorithms have already been approved for colorectal cancer ^27,28^ in Europe, though their widespread adoption is yet to be realized. One possible reason for this delay could be the limited clinical validation against the standard of care ^29^. However, the field of deep learning is progressing rapidly. Over the past year, two techniques have markedly enhanced system performance: transformers and self-supervised learning^30^ (SSL). Furthermore, recent studies have shown that integrating histopathologic imaging with additional modalities, such as genomics, text, clinical imaging, uncovers intermodal relationships and often improves predictive performance ^31–34^.

In this study, we assembled three independent cohorts comprising 6,203 patients with HR+/HER2-EBC with surgically resected primary tumors (**Fig. 1a**). Tissue samples were subjected to H&E staining and immunohistochemical (IHC) analysis for hormone receptors and HER2 according to ASCO/CAP guidelines, and samples were submitted for calculation of RS per clinical practice. For a subset, genomic data from clinical MSK-IMPACT targeted sequencing were also available (**Fig. 1b**). These derivative data were subsequently digitized (**Fig. 1c**) and used for multimodal modeling (**Fig. 1d**).

**Figure 1.**
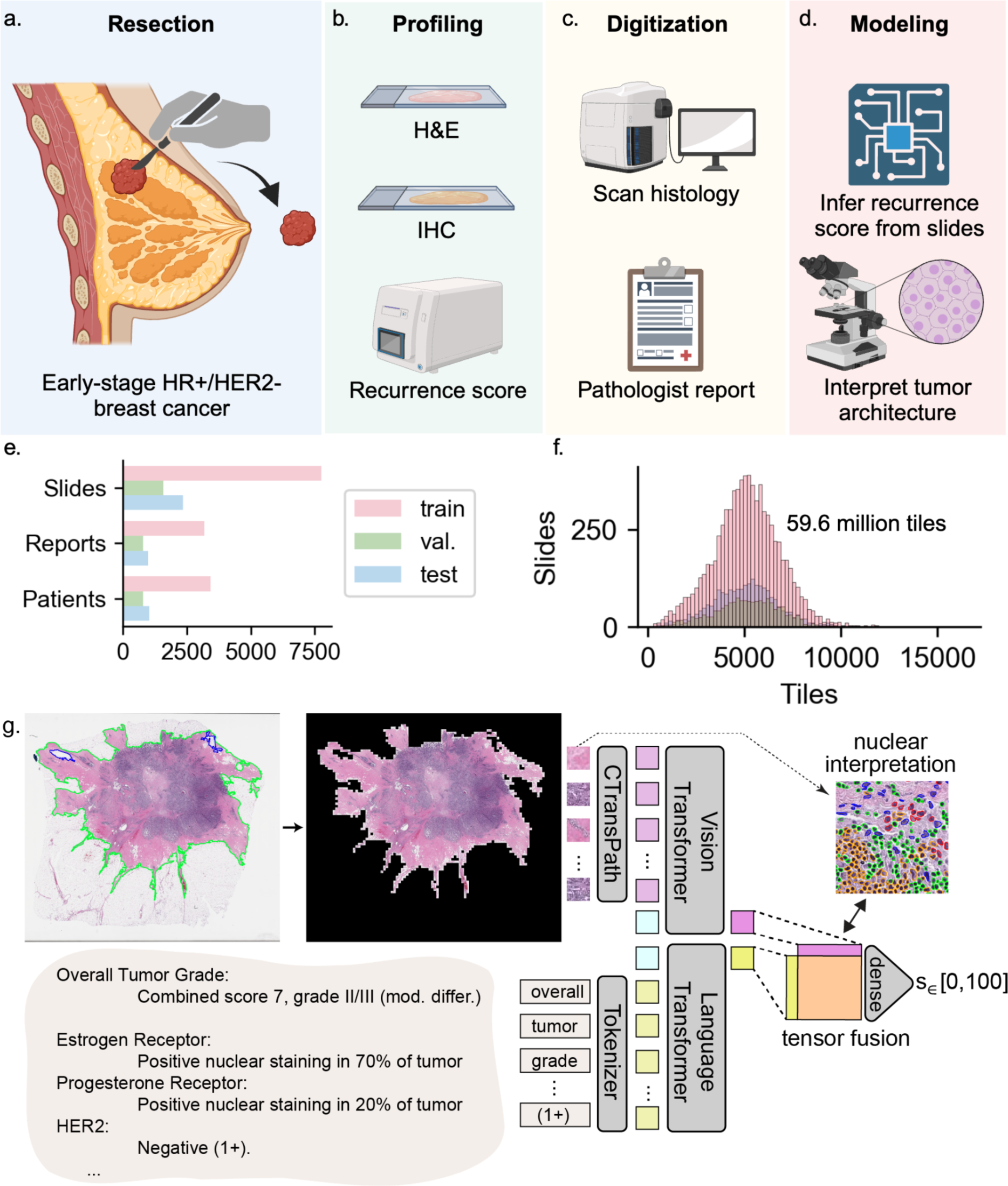
Developing a multimodal transformer model for breast cancer risk. Early-stage breast tumors are (a) resected, (b) profiled histologically (c) digitized, and (d) used for downstream modeling of recurrence risk. (e) Number of pathologic slides, pathology reports, and patients included in each split. (f) Histogram depicting number of slides with a given number of tiles. (g) Tissue detection, tessellation, transformer-based modeling of CTransPath-derived tile embeddings, pathology report scraping, tokenization and transformer-based modeling, nuclear segmentation for interpretation, tensor fusion for multimodal integration. Graphic partially created using BioRender.

We demonstrate that transformer models accurately predict RS from H&E-stained whole-slide images and pathology text reports, and that their integration improves performance beyond that of available nomograms. We also probed the biological interpretability of predictions through computational analysis and suggest clinical operating points to identify high-risk disease. We advance a new model, Orpheus, which has the potential to save testing cost and hasten therapeutic decision-making while maintaining the standard of care based on individual tumor transcriptomes for HR+/HER2-EBC.

## Results

### Data assembly

We curated a retrospective cohort of 5,176 (**Fig. 1e**) patients with HR+/HER2-EBC (MSK-BRCA; **Fig. 1a; Extended Data Fig. 1**) for model training, validation, and testing, whose primary tumors had H&E-stained FFPE tissue specimens available, textual pathology reports, and targeted panel sequencing for a subset (n=331; **Fig. 1b**). We allocated these patients *a priori* into either a withheld test set (20%) or a set used for training and validation (80%; **Supp. Tab. 1**). Moreover, we assembled two additional independent cohorts of whole-slide images derived from patients with HR+/HER2-EBC, IEO-BRCA (452 patients) and MDX-BRCA^35^ (575 patients), for external validation.

### Model training

We developed a transformer model to directly regress the ODX RS from whole-slide images of EBC. To train this architecture, we employed a two-step process. First, we projected each slide’s tissue-containing tiles (**Fig. 1f**) into an informative space using a frozen model trained using SSL on over 30,000 slides (**Fig. 1g**) ^36^. Subsequently, we adapted a transformer architecture ^37^, which was previously validated in a large multicenter study of colorectal cancer ^38^, to map the phenotypic-genotypic correlation between the extracted features and the ODX RS (**Fig. 1g**). The unimodal and multimodal models were trained to regress RS as a continuous variable (**Fig. 1g**).

### Embeddings and predicted score recapitulate clinical and genomic correlates

Uniform manifold approximation and projection (UMAP) over the learned embedding spaces for the visual, linguistic, and multimodal models in the MSK-BRCA test set (**Fig. 2**) revealed that learned embeddings separated somewhat by histologic grade (**Fig. 2a)** and progesterone receptor expression (**Fig. 2b)** in the MSK-BRCA test set (n=1034), with the gradients appearing along a learned, lyre-shaped manifold for the multimodal model. The same was observed for the ODX RS itself (**Fig. 2c**). We further tested the association of predicted scores with genomic features. Limiting to cases with MSK-IMPACT, predicted RS was higher for tumors with *TP53* mutation, *MYC* amplifications, and *PIK3CA* amplifications (**Fig. 2d-f**) and trended slightly higher for specimens with greater fraction of genome altered (**Fig. 2g**).

**Figure 2.**
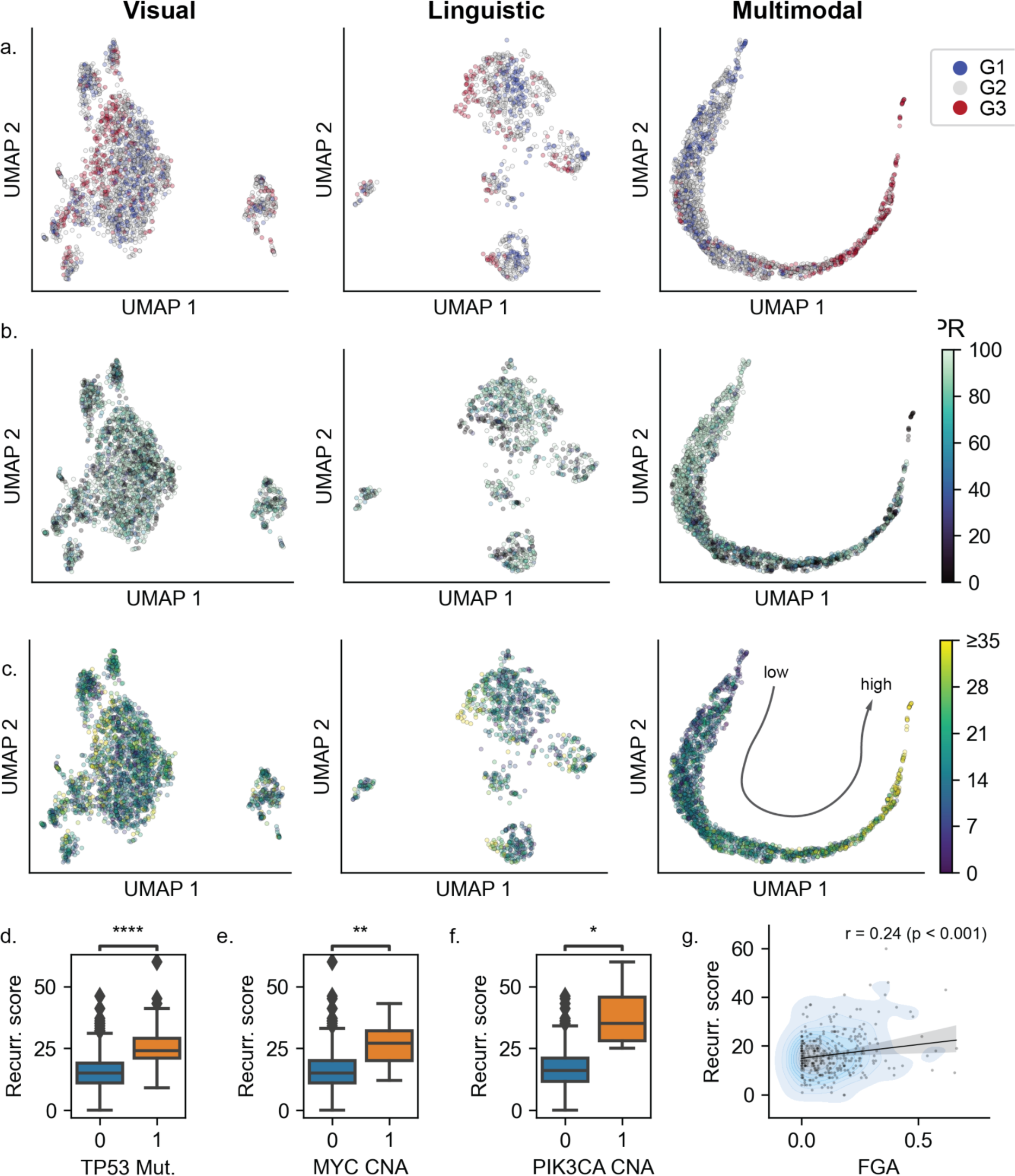
Model distinguishes tumors by biologically meaningful features. UMAP embeddings of the MSK-BRCA test set denoting visual, linguistic, and multimodal representations annotated with (a) histologic grade, (b) progesterone receptor (PR) expression, and (c) recurrence score. (d) *TP53* mutation status, (e) *MYC* copy number amplifications (CNA), *PIK3CA* CNA, and (g) fraction genome altered (FGA) versus recurrence score in the training set. Mann-Whitney U **** *q* < 0.0001, ** *q* < 0.01, * *q* < 0.05.

### Model decisions are clinically explainable

We next sought to interpret the model outputs using attention rollout ^39^. We visualized the last layer’s attention tiles for each slide (**Fig. 3a**), noting that the model designates most tiles as background with low attention scores (**Fig. 3b**). Though the breakdown varies across slides, higher-attention tiles tended to contain invasive and *in situ* carcinoma compared to lower-attention tiles, which are more likely to contain fat and stroma (**Fig. 3c; Extended Data Fig. 2**). The model yielded predicted point-estimate scores alongside 95% confidence intervals (95% C.I.) for use in clinical decision making (**Fig. 3d**). Analogously to the tiles, the importance of word tokens comprising the synoptic pathology report (including fields such as histologic subtype, HR and HER2 IHC staining patterns, histologic grade, anatomic site, and presence of DCIS and LCIS, and other noted histologic features) for the part from which RS was calculated can be analyzed (**Fig. 3e**). Across the whole withheld test set, a word cloud of tokens showed that words around immunohistochemical analyses for estrogen and progesterone receptors and lymphovascular invasion tended to have highest mean relative attention within a report alongside punctuation and descriptions of Nottingham grade (**Fig. 3f**).

**Figure 3.**
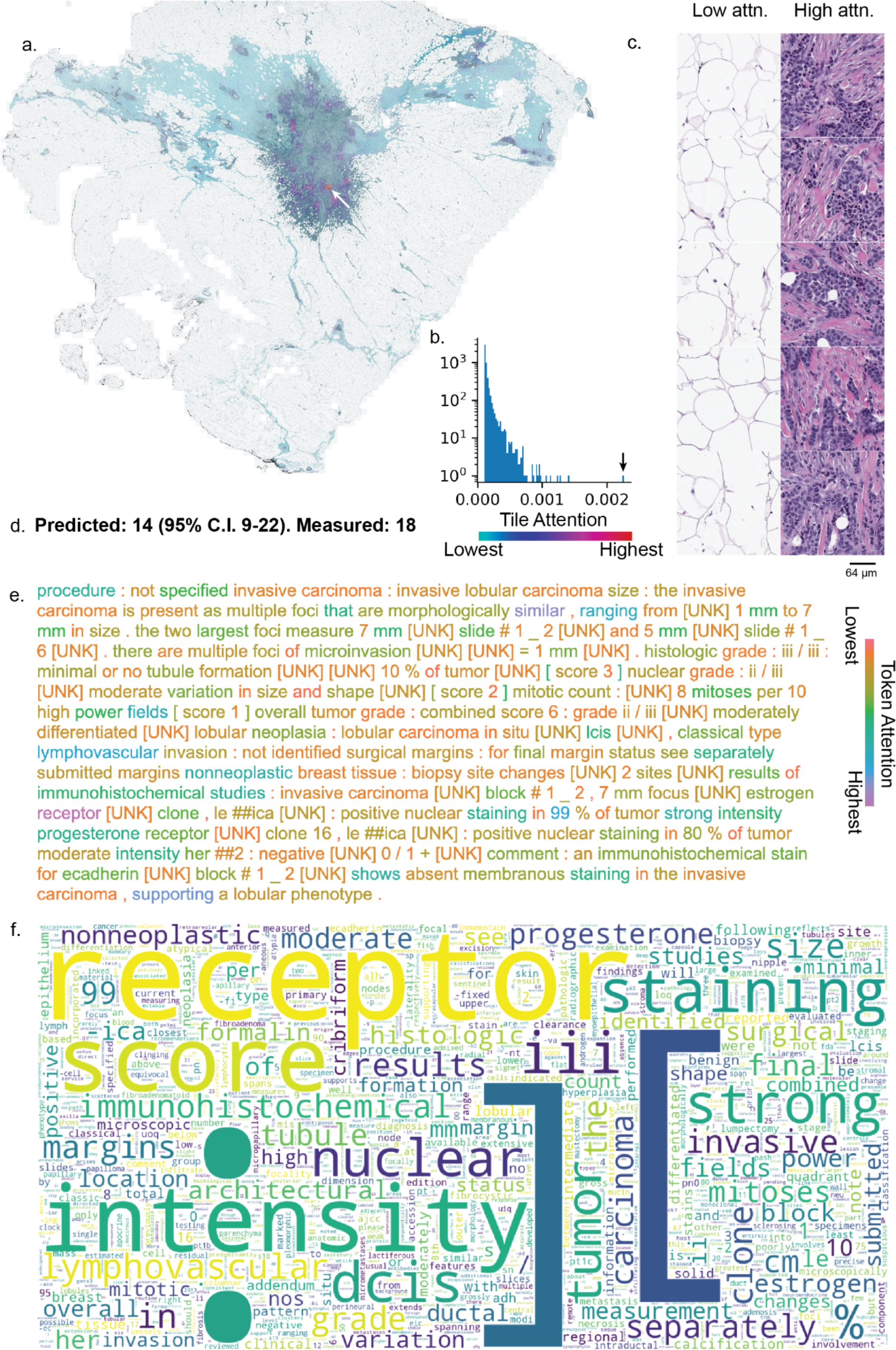
Model decisions are clinically explainable. (a) Foreground tissue colorized by visual attention, plotted in (b). One tile-attention value pair is denoted by the white and black arrows. (c) The five highest- and lowest-attention tiles from (a) at greater magnification. (d) Multimodal prediction with confidence interval and true score. (e) Pathology report-derived tokens colorized by language attention. (f) Whole-cohort token importance cloud with size of word scaled by mean importance across the MSK-BRCA test set. [UNK]: unknown.

### Visual model reproducibly infers recurrence risk

We next tested the reproducibility of the vision model across the three cohorts (**Fig. 4**). In the withheld MSK-BRCA test set, the unimodal whole slide image-based model achieved a Pearson correlation of 0.63 (95% C.I. 0.58 - 0.68, *p* < 10^-4^) and concordance correlation coefficient (CCC) of 0.58 (95% C.I. 0.52 - 0.63; **Fig. 4a**) along with area under the precision-recall curve (AUPRC) of 0.593 (95% C.I. 0.514 - 0.671; **Fig. 4d**) and area under the receiver operating characteristic curve (AUROC) of 0.864 (95% C.I. 0.831 - 0.895; **Fig. 4g**). In the external IEO-BRCA test set, the same model achieved a Pearson correlation of 0.61 (95% C.I. 0.55 - 0.67; p < 10^-4^) and CCC of 0.60 (95% C.I. 0.533 - 0.650; **Fig. 4b**) along with AUPRC of 0.675 (95% C.I. 0.601 - 0.745; **Fig. 4e**) and AUROC of 0.801 (95% C.I. 0.759 - 0.841; **Fig. 4h**). In the external MDX-BRCA test set, which used an inferred, ODX-like RS (see **Methods**), the same model achieved a Pearson correlation of 0.60 (95% C.I. 0.54 - 0.65; p < 10^-4^) and CCC of 0.44 (95% C.I. 0.384 - 0.486; **Fig. 4c**) along with AUPRC of 0.734 (95% C.I. 0.672 - 0.795; **Fig. 4f**) and AUROC of 0.830 (95% C.I. 0.791 - 0.863; **Fig. 4i**). Full results are detailed in the other panels of **Extended Data Fig. 3**, **Fig. 4**, and in **Supp. Tab 2**. In summary, the vision-based model robustly infers RS across three cohorts derived from different medical centers and countries.

**Figure 4.**
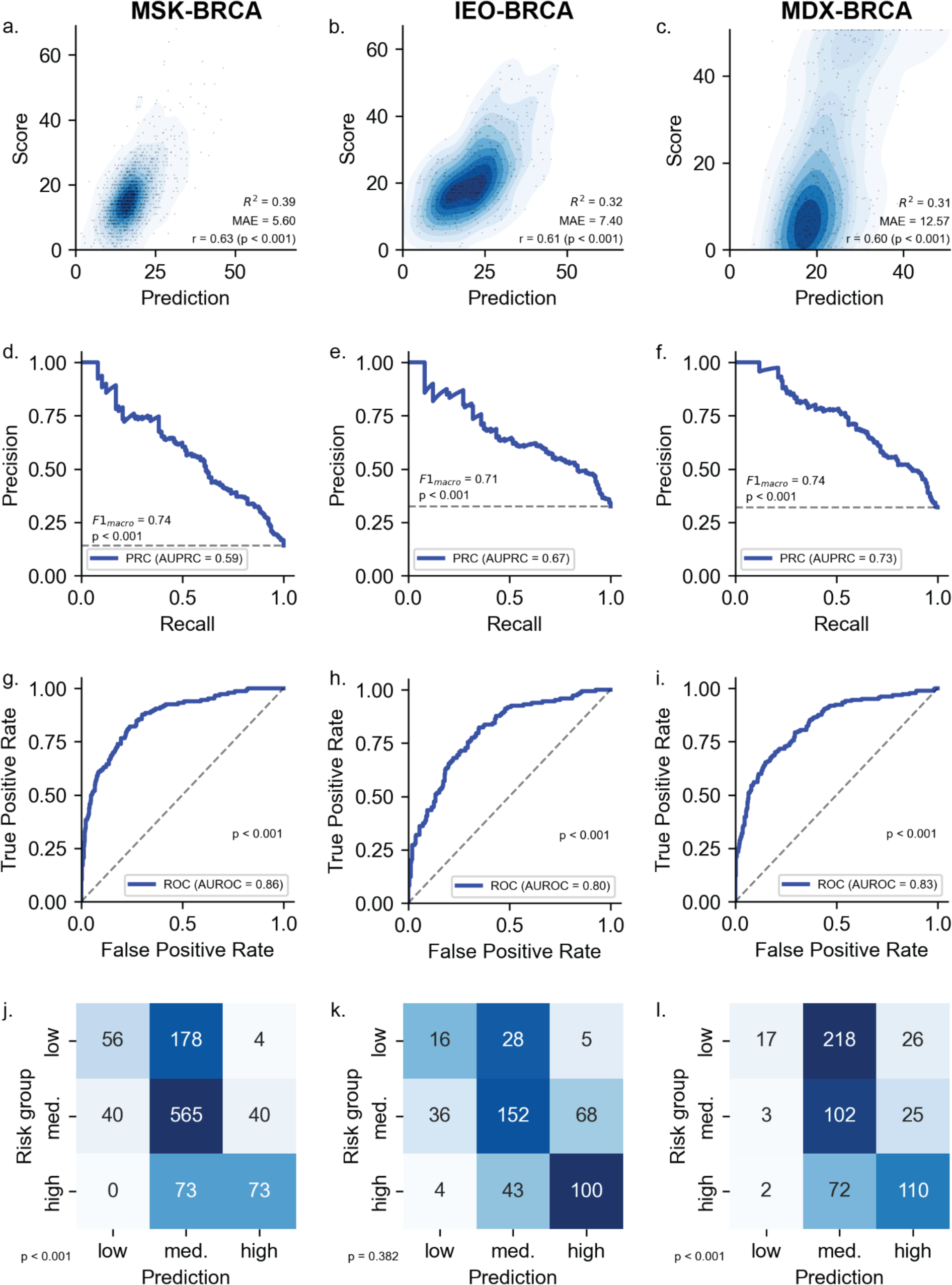
Visual model generalizes internationally to three test cohorts. (a-c) density plots, (d-f) precision-recall curves, (g-i) receiver operating characteristic curves, (j-l) confusion matrices for MSK-BRCA, IEO-BRCA, and MDX-BRCA test sets. MAE: mean absolute error, PRC: precision-recall curve, AUPRC: area under the PRC, AUROC: area under the ROC. *p*-values calculated using (a-c) comparison against the beta distribution, (d-i) 1000-fold permutation testing, (j-l) McNemar’s exact test. Dashed lines in (e-l) represent performance for the minimally informative classifier.

### Model uncovers morphological features

To further explore the model’s capability of correlating histologic features with ODX RS, we identified the most-attended tiles ^39^ for high- and low-risk disease. The nuclei of these tiles were segmented ^40^, and derivative features of cell type proportions and cellular morphology were tabulated (**Fig. 5a**). This revealed a relative abundance of inflammatory cells (**Fig. 5b-c**) and neoplastic cells along with the standard deviation of the neoplastic nuclear area (**Fig. 5d-e**) as some of the features differing significantly between the groups.

**Figure 5.**
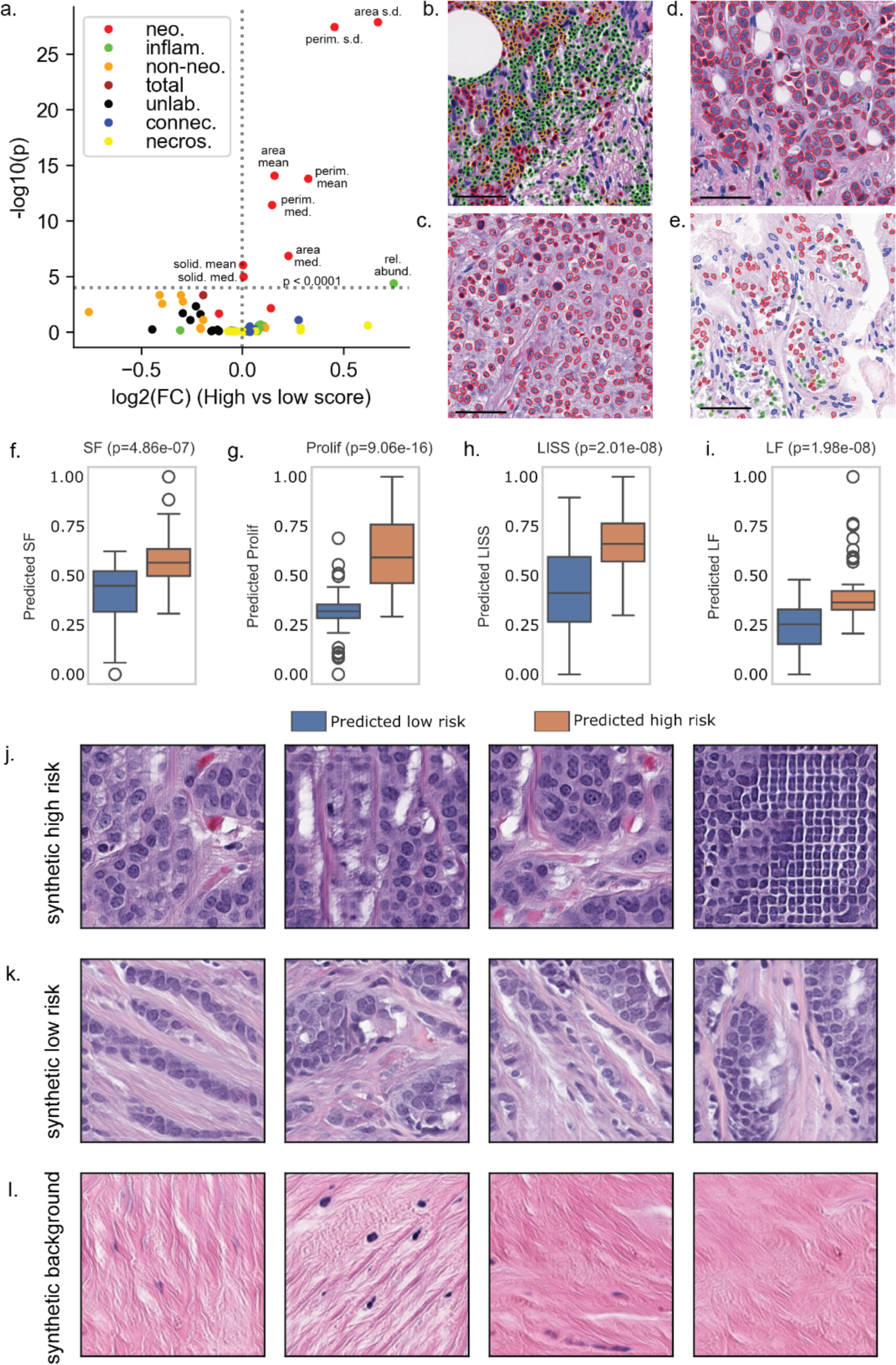
Spatial interpretation of tumors. (a) Association of cellular features with high- and low-risk tissue. (b) High and (c) low relative abundance of inflammatory cells. (d) High and (e) low standard deviation of neoplastic cell area. (f-i) Quantification of stromal fraction (SF), tumor cell proliferation (Prolif), lymphocyte infiltrating signature score (LISS), and lymphocyte fraction (LF) for predicted low- and high-risk patients depicted in blue and orange, respectively, in the MSK-BRCA cohort. *P*-values are generated using an independent t-test (j-l) Depictions of (j) high-risk (including artifact on right), (k) low-risk, and (l) uninformative tissue synthesized by a generative adversarial network. Scale bars denote 64µm. Neo.: neoplastic, inflam.: inflammatory, non-neo.: non-neoplastic, unlab.: unlabeled, connec.: connective, necros.: necrosis

Moreover, a model trained on The Cancer Genome Atlas (TCGA) to infer transcriptomic program activity from imaging features revealed that high-risk disease exhibited greater stromal fraction (p < 10^-4^, n=100) (**Fig. 5f)**, lymphocyte infiltration signature (p < 10^-4^, n=100) (**Fig. 5g)**, tumor cell proliferation (p < 10^-4^, n=100) (**Fig. 5h)**, and leukocyte fraction (p < 10^-4^, n=100) (**Fig. 5i)**. Extending the tumor microenvironment analysis to all three test cohorts corroborated these results, except for the lymphocyte infiltration signature in the MDX cohort which shows no statistically significant difference between the predicted high- and low-risk disease patients (**Extended Data Fig. 4**). As a further study of differences, we also trained a conditional generative adversarial network (GAN) to synthesize fields of view for informative tiles for high- and low-risk disease (**Fig. 5j-l**). Tiles conditioned on the high-risk class depicted confluent clusters of tumor cells with moderate to marked nuclear pleomorphism and prominent nucleoli, and tiles conditioned on the low-risk class depicted trabeculae and clusters of tumor cells with moderate nuclear pleomorphism and inconspicuous nucleoli. Tiles conditioned on the background class depicted stroma without epithelial cells.

### Integrating imaging and language information improves stratification

In the MSK-BRCA test set, the unimodal text report-based model achieved a Pearson correlation of 0.58 (95% C.I. 0.51 - 0.64, *p* < 10^-4^) and CCC of 0.55 (95% C.I. 0.478 - 0.606; **Extended Data Fig. 5c**) along with AUPRC of 0.539 (95% C.I. 0.455 - 0.628; **Extended Data Fig. 5f**) and AUROC of 0.820 (95% C.I. 0.779 - 0.854; **Extended Data Fig. 5i**). Full results are detailed in the other panels of **Extended Data Fig. 5** and in **Supp. Tab. 2**.

We then tested if multimodal integration could improve on the image or text models alone, using tensor fusion ^41^ of the transformer-based embeddings. In the MSK-BRCA test set, the full multimodal model achieved a Pearson correlation of 0.68 (**Fig. 6a**; 95% C.I. 0.64-0.73, *p* < 10^-4^) and CCC of 0.65 (95% C.I. 0.59-0.70). For classification of high-risk (RS ≥ 26) disease, the AUPRC was 0.64 (*p* < 10^-4^; 95% C.I. 0.56 - 0.71), with a macro-averaged *F1* score of 0.75 (**Fig. 6b**). The CCC and Pearson’s correlation based on multimodal scores were higher than those based on unimodal scores (**Fig. 6c,f**). AUROC was 0.88 (**Fig. 6d**; 95% C.I. 0.85 - 0.91, *p* < 10^-4^). Using <12 and >25 as thresholds for low-, intermediate-, and high-risk disease, the confusion matrix for the withheld test set is depicted in **Fig. 6e**, showing very low confusion between the extrema and moderate confusion between intermediate and extreme categories (*p* < 10^-4^).

**Figure 6.**
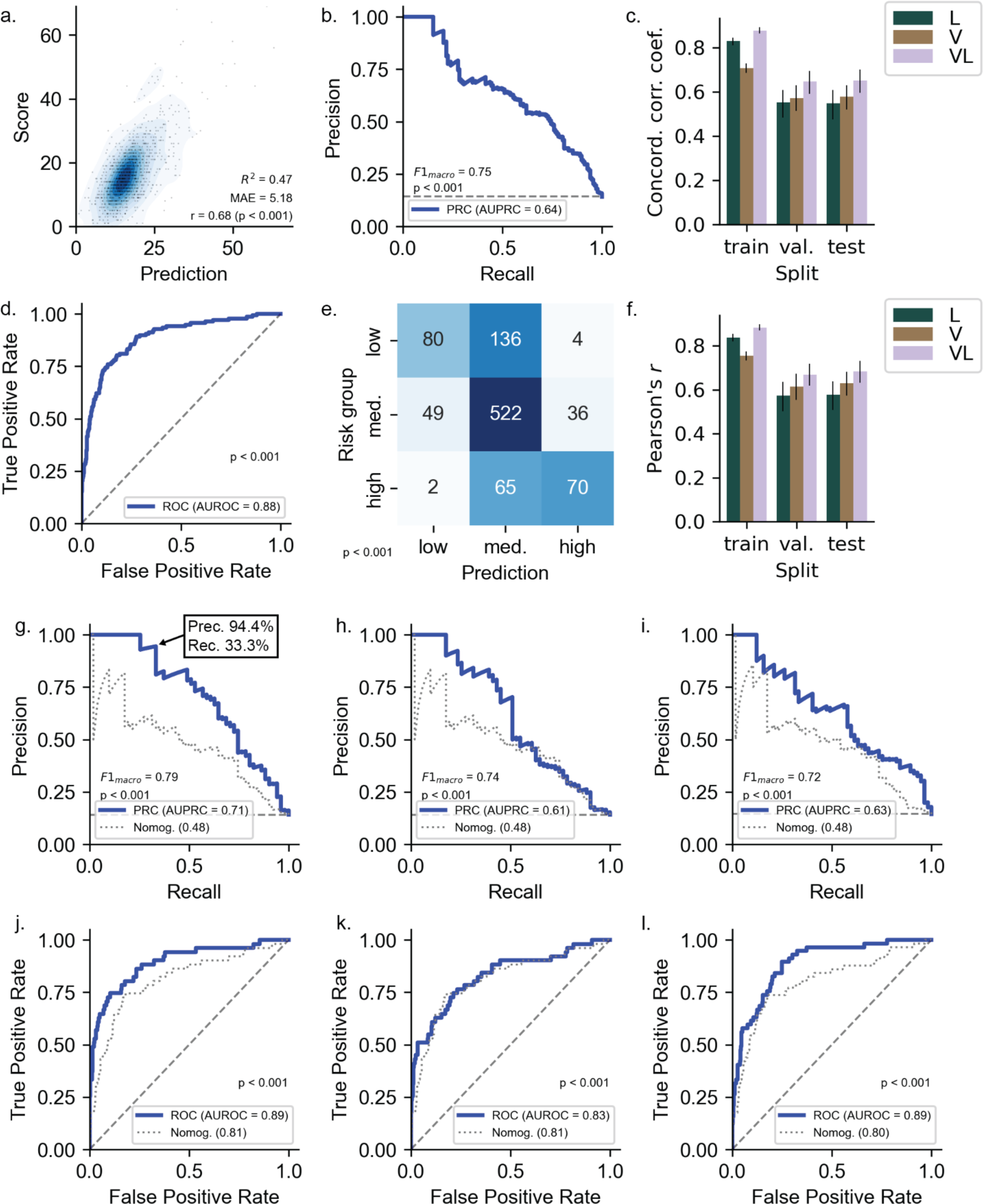
Multimodal model performance and benchmarking in the MSK-BRCA test set, (a) regression of predicted versus true recurrence scores, (b) precision-recall curve for high-risk disease, (c) concordance correlation coefficient for all data splits and models, (d) receiver operating characteristic curve for high-risk disease (e) confusion matrix using score cutoffs of 11 and 25, (f) Pearson correlation for all data splits and models. (g-i) PRCs and (j-l) ROCs for multimodal, language, and vision models, respectively, compared against a clinical nomogram in the full information setting. MAE: mean absolute error, PRC: precision-recall curve, AUPRC: area under the PRC, L: language model, V: vision model, VL: multimodal model, ROC: receiver operating characteristic curve, AUROC: area under the ROC. Error bars (c,f) and shaded linear uncertainty in (a) represent 95% confidence intervals by bootstrapping. *p*-values calculated using (a) comparison against the beta distribution, (b,d) 1000-fold permutation testing, (e) McNemar’s exact test. Dashed lines in (a,d) represent performance for the minimally informative classifier.

Next, we analyzed the subset of the MSK-BRCA test set with available tumor grades and IHC-derived HR status in the text report as extracted by regular expressions (those without matches by regular expressions were excluded). For this set, we compared the ability to discriminate high-risk disease of a nomogram based on clinical and pathologist-annotated features ^17^ to that of the multimodal (**Fig. 6g**), text-based (**Fig. 6h**), and image-based (**Fig. 6i**) models. The multimodal model achieved an AUROC of 0.89 and AUPRC of 0.71 (95% C.I. 0.60 - 0.82), the vision model achieved an AUPRC of 0.63 (95% C.I. 0.50 - 0.75), and the language model achieved an AUPRC of 0.61 (95% C.I. 0.48 - 0.73). By comparison, the nomogram ^17^ achieved an AUPRC of 0.49 (95% C.I. 0.36 - 0.64). For the multimodal model, we suggest an operating point of 29.8 with 94.4% precision and 33.3% recall (**Fig. 6g**).

### Assessing clinical utility as a triaging tool for low- and high-risk disease

We next tested the utility of our multimodal transformer model as a pre-screening tool to reduce the load of laboratory testing for breast cancer recurrence in clinical workflows. Performing a sensitivity analysis, we manually selected a threshold in the test set of the MSK-BRCA (n=2338) cohort which yields the highest sensitivity for the largest percentage of the cohort’s population. This resulted in a sensitivity of 0.93 for 34% of the population with a threshold of < 16 for the predicted recurrence risk score to determine intermediate/low-risk patients (**Extended Data Fig. 6a**) for the test set of the MSK-BRCA cohort. Applying this threshold on the IEO-BRCA (n=452) and MDX-BRCA (n=572) cohorts, we achieved a sensitivity of 0.94 for 25% (**Extended Data Fig. 6b**) and 0.96 for 18% of the populations (**Extended Data Fig. 6c**), respectively.

Similarly, we conducted a specificity analysis, wherein we manually selected a threshold in the MSK-BRCA test set to yield the highest specificity for the largest percentage of the cohort’s population. This resulted in a specificity of 0.93 for 13% of the population with a threshold of > 25 for the predicted RS to identify high-risk patients (**Extended Data Fig. 6d**) for the test set of the MSK-BRCA cohort. Applying this threshold on the subsequent cohorts, we achieved a specificity of 0.76 for 40% of the population (**Extended Data Fig. 6e**) and 0.85 for 31% of the population (**Extended Data Fig. 6f**) in the IEO-BRCA and MDX-BRCA cohorts, respectively. We repeated the analyses stratified by age and nodal status, specifically patients with node-negative disease below 50 years of age (**Extended Data Fig. 7**) and patients with 1-3 positive nodes above or equal to 50 years of age (**Extended Data Fig. 8**), with similar performance metrics in all cohorts regardless of age and nodal status.

In summary, the Orpheus model has the potential to accurately and highly confidently identify patients with high-risk disease. In a potential use case, adjuvant chemotherapy could be recommended for a selected subset of high-confidence high-risk patients without multigene assay testing (**Extended Data Fig. 9**).

## Discussion

Proper selection of patients with HR+/HER2-EBC who can safely omit adjuvant chemotherapy is a priority in clinical practice. Validated multigene assays, such as ODX RS, have the power to tailor adjuvant treatment-decision making in this setting. However, due to fiscal and logistical barriers, they have faced limited adoption in non-American healthcare systems despite long-standing recommendations for their use ^13^. In this study, we show in a large-scale analysis comprising thousands of patients with HR+/HER2-EBC from internationally distinct cohorts that machine learning on whole-slide images accurately and reproducibly infers RS from routinely available H&E-stained specimens or their corresponding text report.

These models and their multimodal combination outperform a nomogram ^17^ using clinico-pathologic features, such as IHC-derived progesterone/estrogen receptor positivity, tumor size, lobular versus ductal histology, Nottingham grade, and clinical features, such as age ^17^. The optimal operating point accurately retrieves one third of high-risk disease with minimal false positives, potentially enabling physicians to forgo testing on one in three newly diagnosed patients. If deployed clinically, the improved accuracy of this technique and reduced requirement for manual curation would be expected to further improve the cost effectiveness ^16^. Few institutions have fully digital pathology workflows, but commercial services offer scanning for USD 35 per slide (Biochain, Newark, CA), and model inference is relatively inexpensive at USD 0.90 per hour (Amazon Web Services, Seattle, WA), with average model inference requiring significantly less than one minute per slide. Assuming a cost of USD 4,000 per slide for the laboratory assay ^14,15^, a hypothetical fee of USD 50 per slide for the artificial intelligence-derived test, and our empirically estimated recall of 33.3% (where any patients with scores below the operating point are sent for laboratory assay measurement), this results in an estimated average savings of USD 1,271 per patient without compromising the standard of precision oncology. Moreover, the speed of this assay could enable novel uses such as more precisely defining populations that will benefit from the use of neoadjuvant therapies beyond the currently used clinical characteristics and without the requirement for additional biopsies ^42^.

Further analysis of the proposed method as a potential pre-screening tool revealed consistent sensitivity and specificity to identify patients with high- and low-risk tumors for approximately 50% of the population across three distinct large cohorts. Notably, the risk prediction results remained generally unaffected by subgroup analyses taking into account age and nodal status, despite being clinicopathological factors which impact the recurrence risk in patients with breast cancer, and therefore influence the risk category threshold ^43–45^.

With proper regulatory approval, adoption of clinical artificial intelligence in this paradigm—as a triaging or support tool—is a more measured approach than outright replacement of genomic tests or physician judgment and is more likely to result in widespread adoption. This computational tool has the added benefit of providing confidence intervals rather than pure point estimates, enabling integration of uncertainty into clinical decision making. Similarly, an additional benefit over prior nomograms is the provision of a continuous recurrence score rather than mere risk category, enabling downstream use of the RS for emerging uses, such as the patient selection for adjuvant radiotherapy after breast-conserving surgery ^46,47^, for neoadjuvant chemotherapy and in selecting patients for clinical trials. The architecture makes use of self-supervised learning to enable training on under 5,000 patients and Cartesian product with dimensionality reduction ^41^ to commingle the text-based and image-based features.

For each specimen, the model also generates an annotated report of the text and image used to estimate the recurrence score. Though deep learning suffers from a general lack of interpretability ^48^, the attention paid to each token in the text or each tile in the image enables ordering physicians to perform qualitative quality controls. In the analysis of word importance, “lymphovascular” and “invasion” also appeared, reflecting the association of lymphovascular invasion with disease recurrence risk ^49^, though this is not a feature of the clinicopathologic nomogram ^17^. We furthermore saw colocalization of lower progesterone receptor percent positivity and higher Nottingham grade with higher recurrence score in the models’ learned embedding spaces, both of which are known associations. Informative imaging tiles tended to contain invasive carcinoma but sometimes also contained stroma and other known correlates of recurrence risk, such as lobular carcinoma *in situ* ^50^.

We also sought to identify quantitative biologic features underpinning high- and low-risk disease in tiles deemed informative by the model, including cell-level analyses that have begun to yield fruit in other studies ^31,33,51^. We found that abundance of inflammatory cells corresponded with higher-risk disease, corroborating prior studies that tumor infiltrating lymphocytes are a negative prognostic factor in HR+/HER2-EBC and are associated with a somewhat higher RS ^52–55^. The association is widely validated^56^, but its nature is unknown. Based on our findings that both the fraction of genome altered and inflammatory cell infiltrate are higher for high-risk disease, we submit that one putative mechanism is that more aggressive disease exhibits greater chromosomal instability, which in turn increases intratumoral inflammation. The greater nuclear pleomorphism and nucleolar prominence in high-risk tumors synthesized by the GAN associates with higher Nottingham grade ^57^. Our cohort also recapitulated previously uncovered genomic biomarkers with adverse prognostic implications, namely *MYC* amplification^58^*, PIK3CA* amplification^59^, and *TP53* mutation^60^. By estimating transcriptomic programs from images using our validated model ^61^, proliferation was also found to be higher in our analysis of patients with predicted high risk, correlating with grade, the *MKI67* gene included in the calculation of the RS, and perhaps explaining the empiric association of more heterogeneous areas and perimeters of cancer cells with higher risk disease. The elevated presence of the lymphocyte infiltrating signature score and leukocyte fraction in predicted high-risk tumors hints at biologically aggressive cancer and was found to positively correlate with recurrence risk of HR+/HER2-EBC ^62,63^. Furthermore, our finding of increased stromal fraction in predicted tumors with predicted high risk across all cohorts corroborates the association between high stromal fraction and cancer-associated fibroblasts ^56^ with worse prognosis in various breast cancer subtypes ^64,65^, with our analysis specifically building a case for HR+/HER2-tumors. The finding of stroma in the background tiles generated by the GAN is possibly due to the prevalence of uninformative stromal areas further from the tumor-stroma interface. Together, these findings show that our new deep learning method can be used as a tool to make biological discoveries and suggest mechanistic hypotheses.

The greatest limitation of this study is that it relies on a laboratory assay–albeit rigorously validated–for ground truth rather than clinical outcomes, and thus the model cannot be detected to discriminate risk of distal recurrence better than ODX RS. That is, using the RS as the ground truth will penalize deviations, even if they could hypothetically be more closely associated with true clinical risk of recurrence. As clinical outcomes become more readily available at scale, we aim to test the ability of such models on censored time-to-event modeling, in this case for distal recurrence. A second limitation is the reliance on deep transformer architectures: though they are ensconced as the workhorse of modern artificial intelligence^66^, they lack true interpretability, with *post hoc* explainability instead standing in^48^. That is, we must deploy methods to interpret the model’s decisions and robustness rather than asking the model to directly explain its reasoning. Nonetheless, recurrence risk models with inherit interpretability capabilities tend to perform worse on the main evaluation metrics such as the AUROC^67^.

In summary, we have developed Orpheus, an artificial intelligence model that accurately infers Oncotype DX ® Recurrence Score from H&E-stained whole slide images, and validated it across three independent cohorts totalling 6,203 patients internationally. We have rigorously analyzed the biological and clinical underpinnings of our model’s decisions and suggest that our architecture can be tailored for application to rapid biomarker inference in any tumor type.

## Code availability

All source code is available under an open-source license on GitHub. The multimodal modeling package, Orpheus, is available at https://github.com/kmboehm/orpheus. The pre-processing pipeline for whole-slide images is found at https://github.com/KatherLab/STAMP, and our code for regressing transcriptomic programs from images is found at https://github.com/KatherLab/marugoto/releases/tag/v1.0.0-regression. The GAN was trained using https://github.com/POSTECH-CVLab/PyTorch-StudioGAN, with our weights and configuration parameters at https://www.synapse.org/breastGAN. The code to calculate nuclear features based on HoverNet inference is at https://gist.github.com/kmboehm/aea77f24a9cdbb1f246dacaae812053d.

---

**Extended Data Fig 1:**
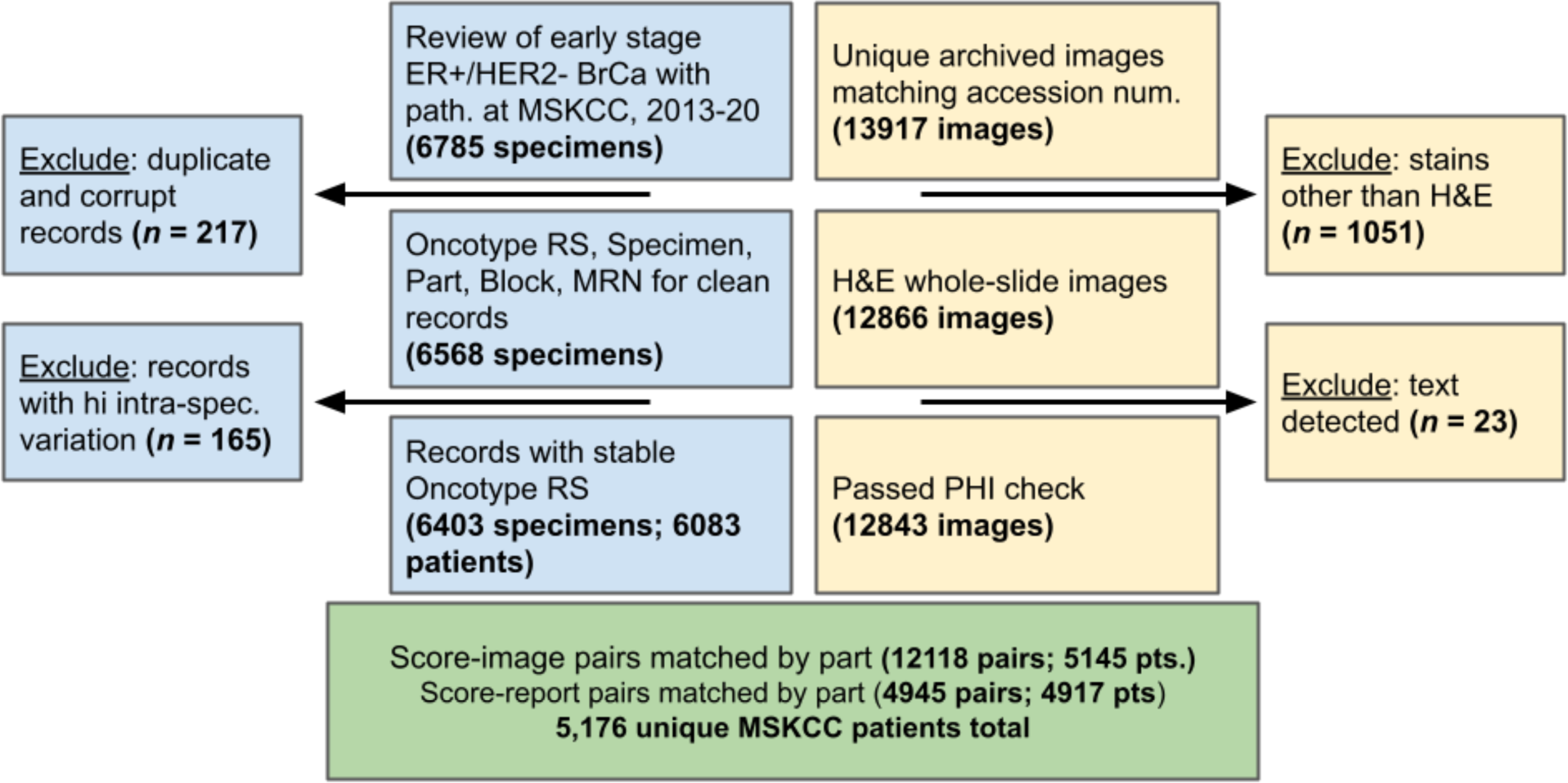
Case inclusion diagram. Depicts slides (yellow) and patients (blue) joined to form the full cohort of paired slides and reports with recurrence scores (green).

**Extended Data Fig 2:**
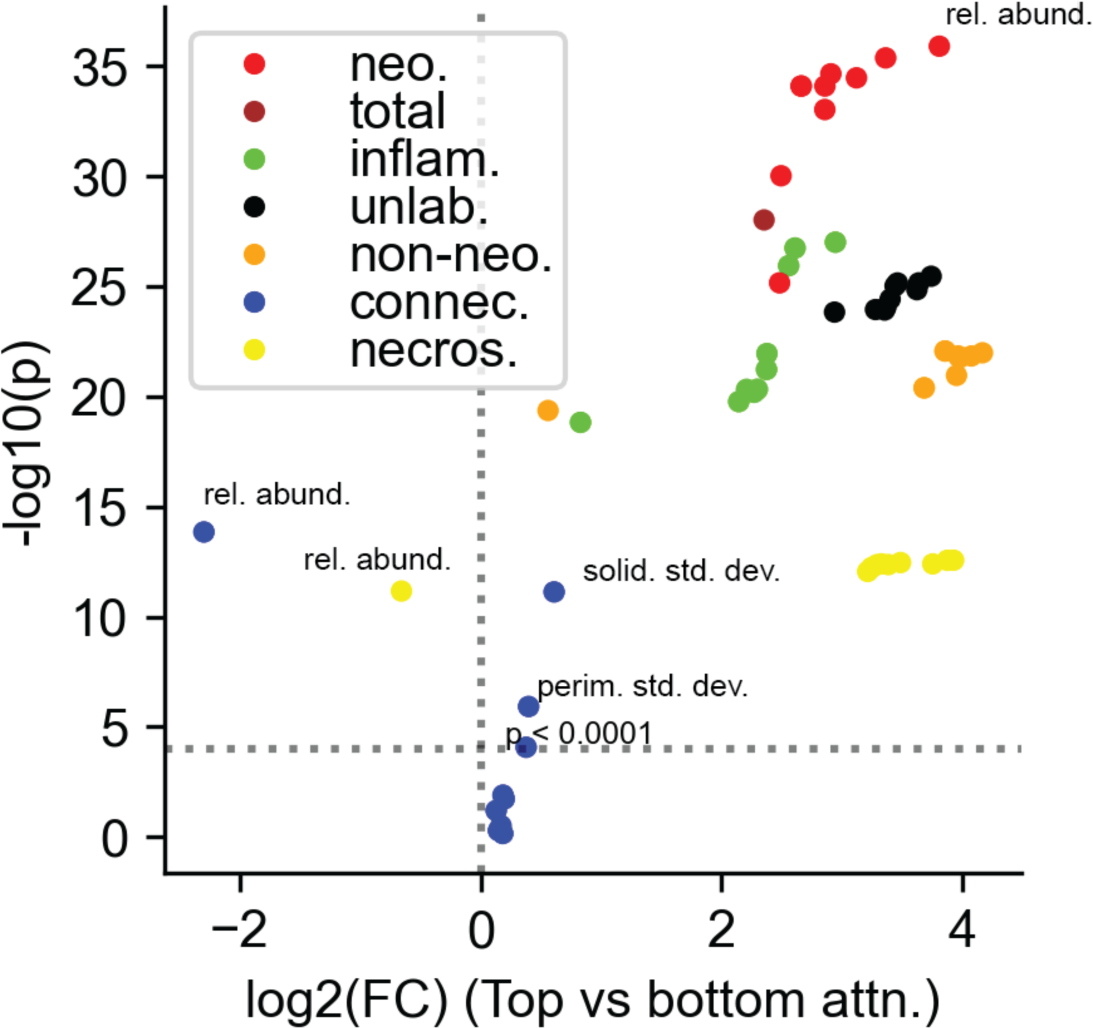
Quantitative analysis of high-versus low-attention tiles. Full feature titles available in source data.

**Extended Data Fig 3.**
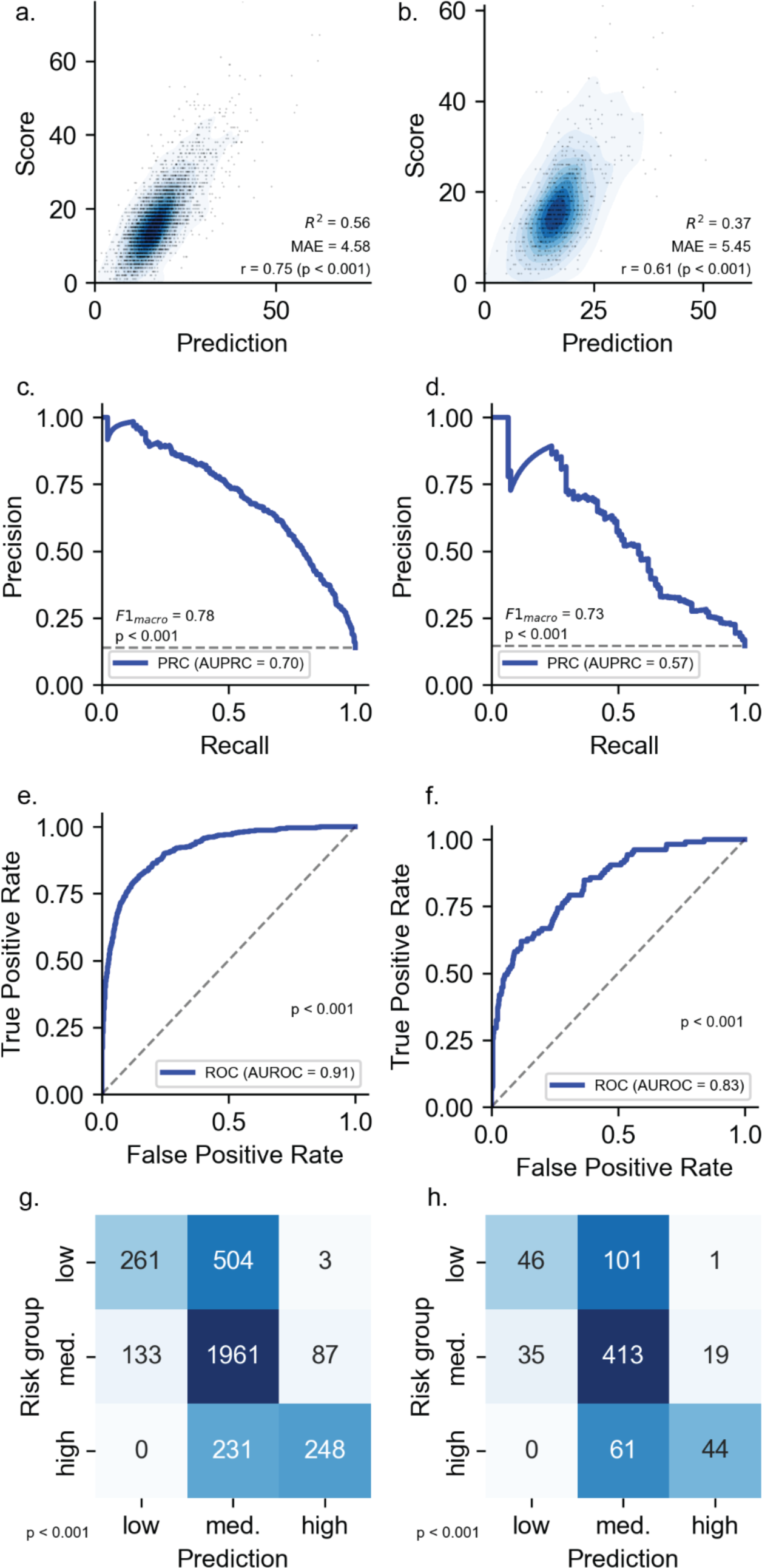
MSK-BRCA training and validation unimodal vision model performance. (a-b) density plots, (c-d) precision-recall curves, (e-f) receiver operating characteristic curves, (g-h) confusion matrices for MSK-BRCA training and validation sets (left to right). MAE: mean absolute error, PRC: precision-recall curve, AUPRC: area under the PRC, AUROC: area under the ROC. *p*-values calculated using (a-d) comparison against the beta distribution, (e-l) 1000-fold permutation testing, (m-p) McNemar’s exact test. Dashed lines in (e-l) represent performance for the minimally informative classifier.

**Extended Data Fig 4:**
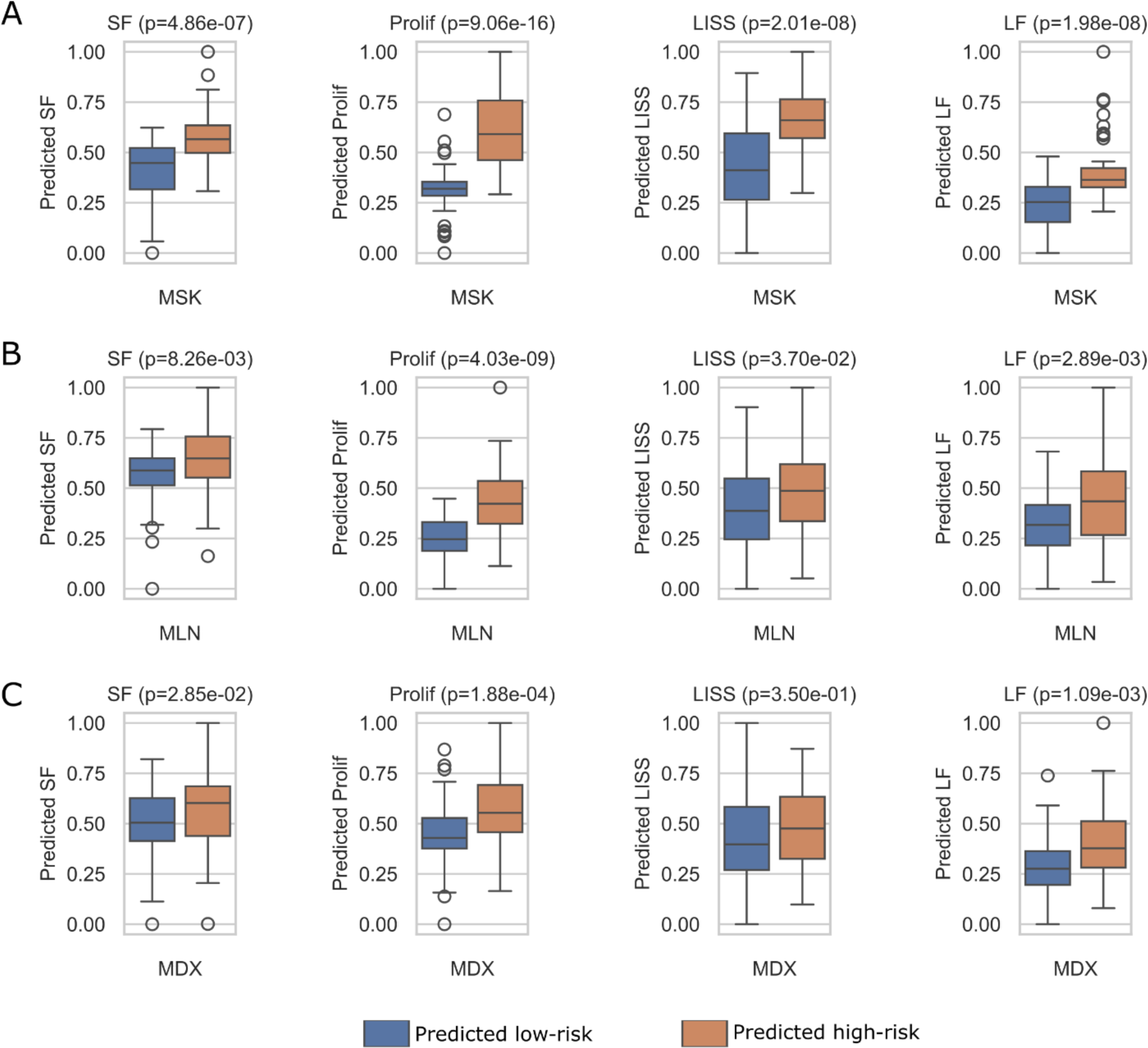
Tumor microenvironment quantification of the top attention tiles of the recurrence risk vision prediction model. Quantification of the tumor microenvironment for the top 50 predicted high- and low-risk patients by the recurrence risk vision prediction model, specifically for the stromal fraction (SF) and leukocyte fraction (LF) as assessed via DNA methylation analysis, lymphocyte infiltrating signature score (LISS) and proliferation (Prolif) as measured by RNA expression for the **a.** MSK cohort (n=100), **b.** IEO cohort (n=100) and **c.** MDX cohort (n=100). Statistical significance is measured by an independent t-test, indicating a difference in sample means between predicted high- and low-risk patients (p < 0.05).

**Extended Data Fig 5.**
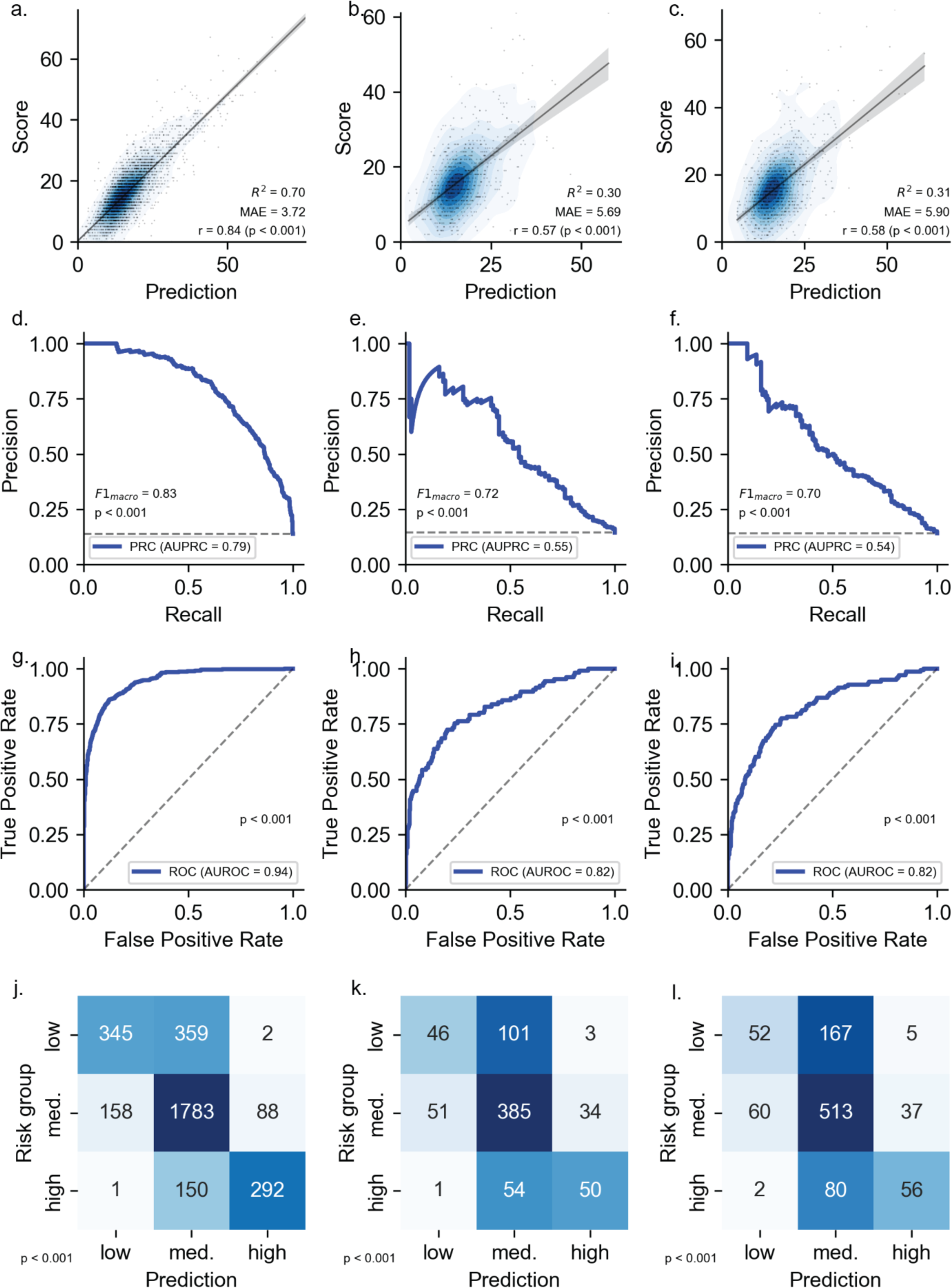
Unimodal language model performance. (a-c) density plots, (d-f) precision-recall curves, (g-i) receiver operating characteristic curves, (j-l) confusion matrices for training, validation, and MSKCC test sets (left to right). MAE: mean absolute error, PRC: precision-recall curve, AUPRC: area under the PRC, AUROC: area under the ROC. *p*-values calculated using (a-d) comparison against the beta distribution, (e-l) 1000-fold permutation testing, (m-p) McNemar’s exact test. Dashed lines in (e-l) represent performance for the minimally informative classifier.

**Extended Data Fig. 6:**
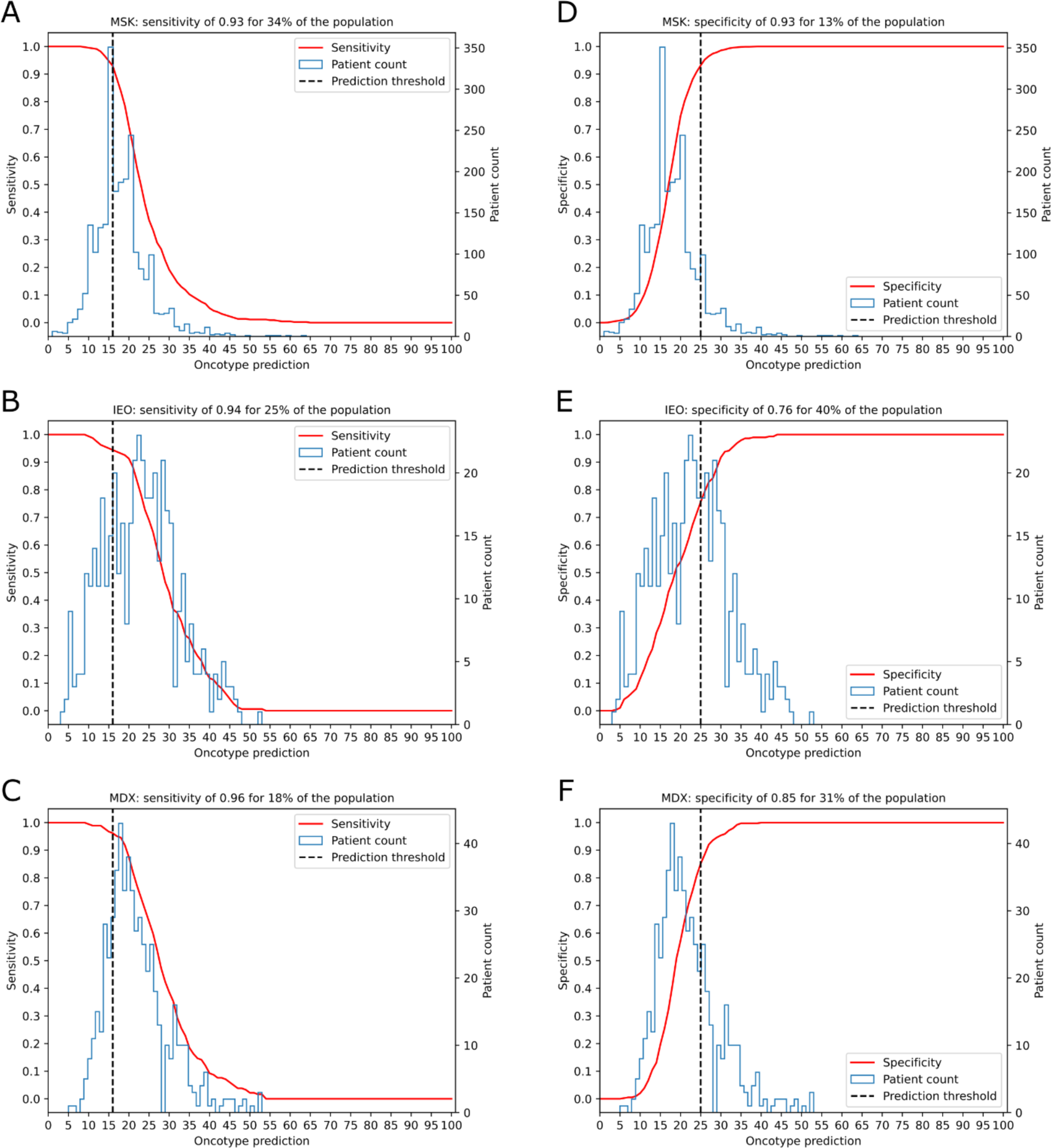
Sensitivity and specificity analysis for the predicted recurrence risk scores of all patients. The sensitivity and specificity are calculated using a threshold for the predicted recurrence risk score of < 16 and ≥ 25, respectively. The thresholds are determined in the MSK cohort (n=2338) and are set in the external IEO (n=452) and MDX (n=572) cohorts. The analysis does not account for age and nodal status. The sensitivity versus the patient count is plotted for the **a.** MSK, **b.** IEO and **c.** MDX cohorts. Moreover, the specificity versus the patient count is plotted for the **d.** MSK, **e.** IEO and **f.** MDX cohorts.

**Extended Data Fig. 7:**
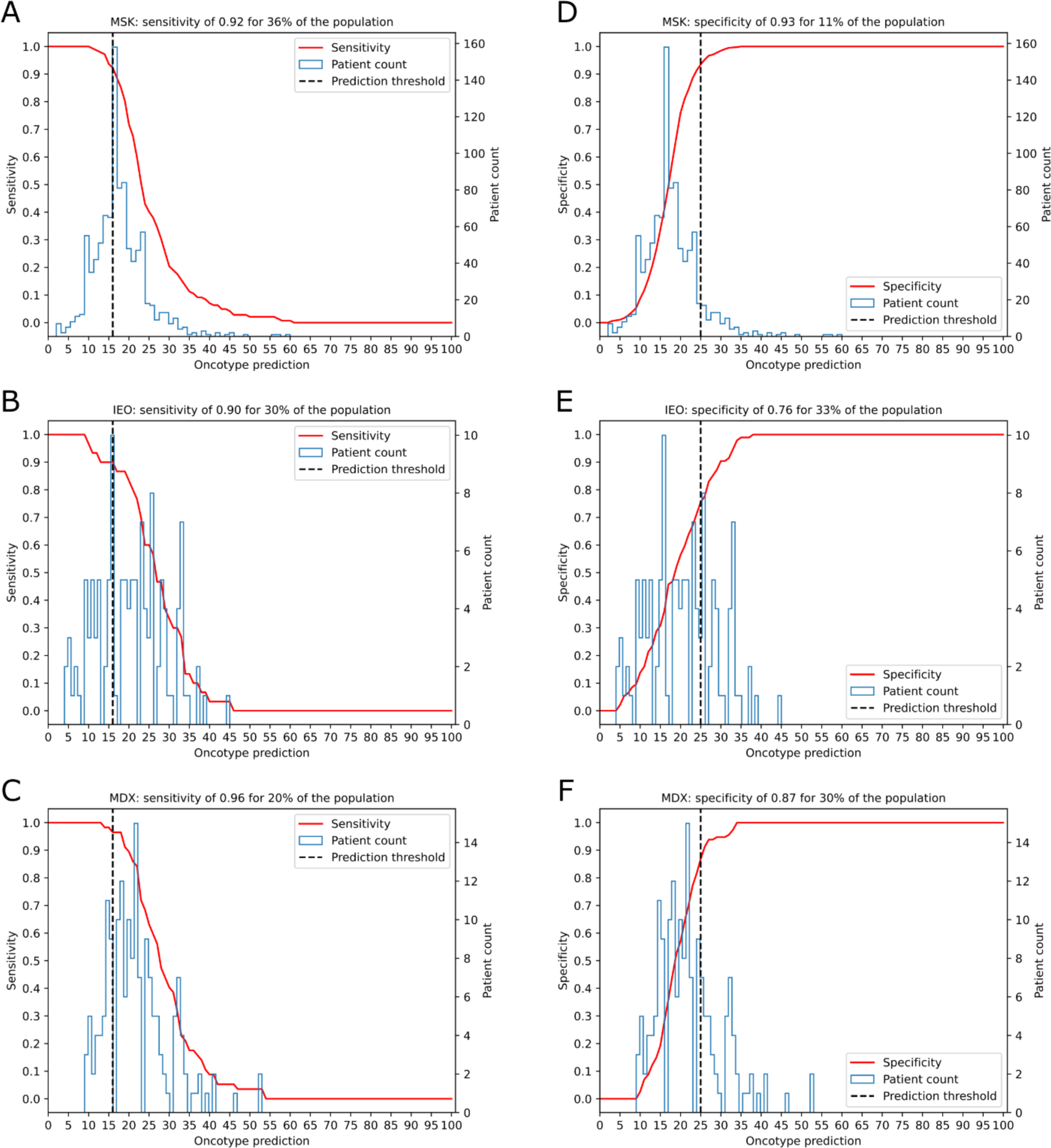
Sensitivity and specificity analysis for the predicted recurrence risk scores of all patients above 50 years of age and having 1-3 positive nodes. The sensitivity and specificity are calculated using a threshold for the predicted recurrence risk score of < 16 and ≥ 25, respectively. The thresholds are determined in the MSK cohort (n=987) and are set in the external IEO (n=124) and MDX (n=171) cohorts. The analysis is focussed on the patient subset above 50 years of age and having 1-3 positive nodes. The sensitivity versus the patient count is plotted for the **a.** MSK, **b.** IEO and **c.** MDX cohorts. Moreover, the specificity versus the patient count is plotted for the **d.** MSK, **e.** IEO and **f.** MDX cohorts.

**Extended Data Fig. 8:**
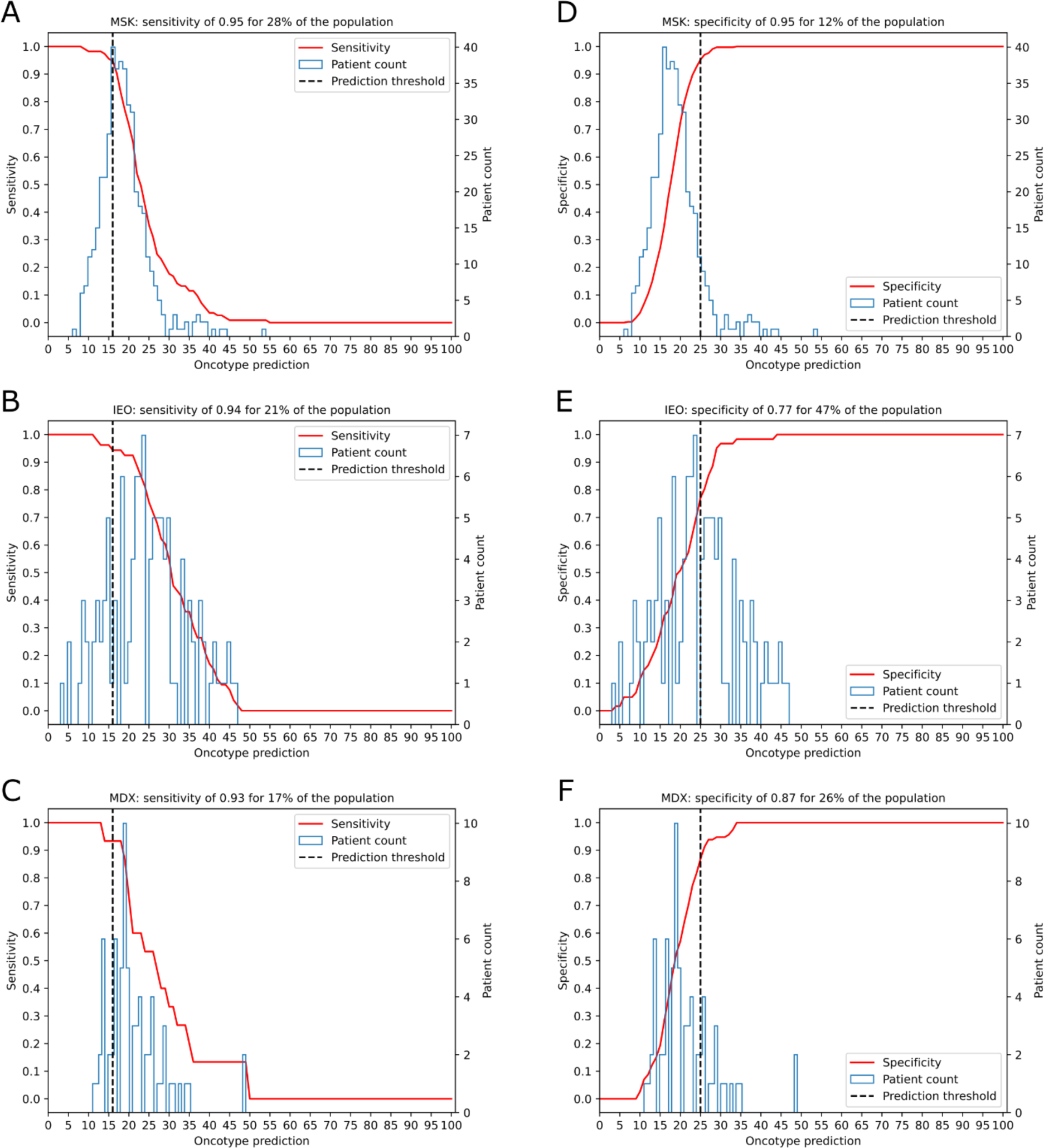
Sensitivity and specificity analysis for the predicted recurrence risk scores of all patients below 50 years of age and 0 positive nodes. The sensitivity and specificity are calculated using a threshold for the predicted recurrence risk score of < 16 and ≥ 25, respectively. The thresholds are determined in the MSK cohort (n=450) and are set in the external IEO (n=114) and MDX (n=70) cohorts. The analysis is focussed on the patient subset below 50 years of age and having 0 positive nodes. The sensitivity versus the patient count is plotted for the **a.** MSK, **b.** IEO and **c.** MDX cohorts. Moreover, the specificity versus the patient count is plotted for the **d.** MSK, **e.** IEO and **f.** MDX cohorts.

**Extended Data Fig. 9:**
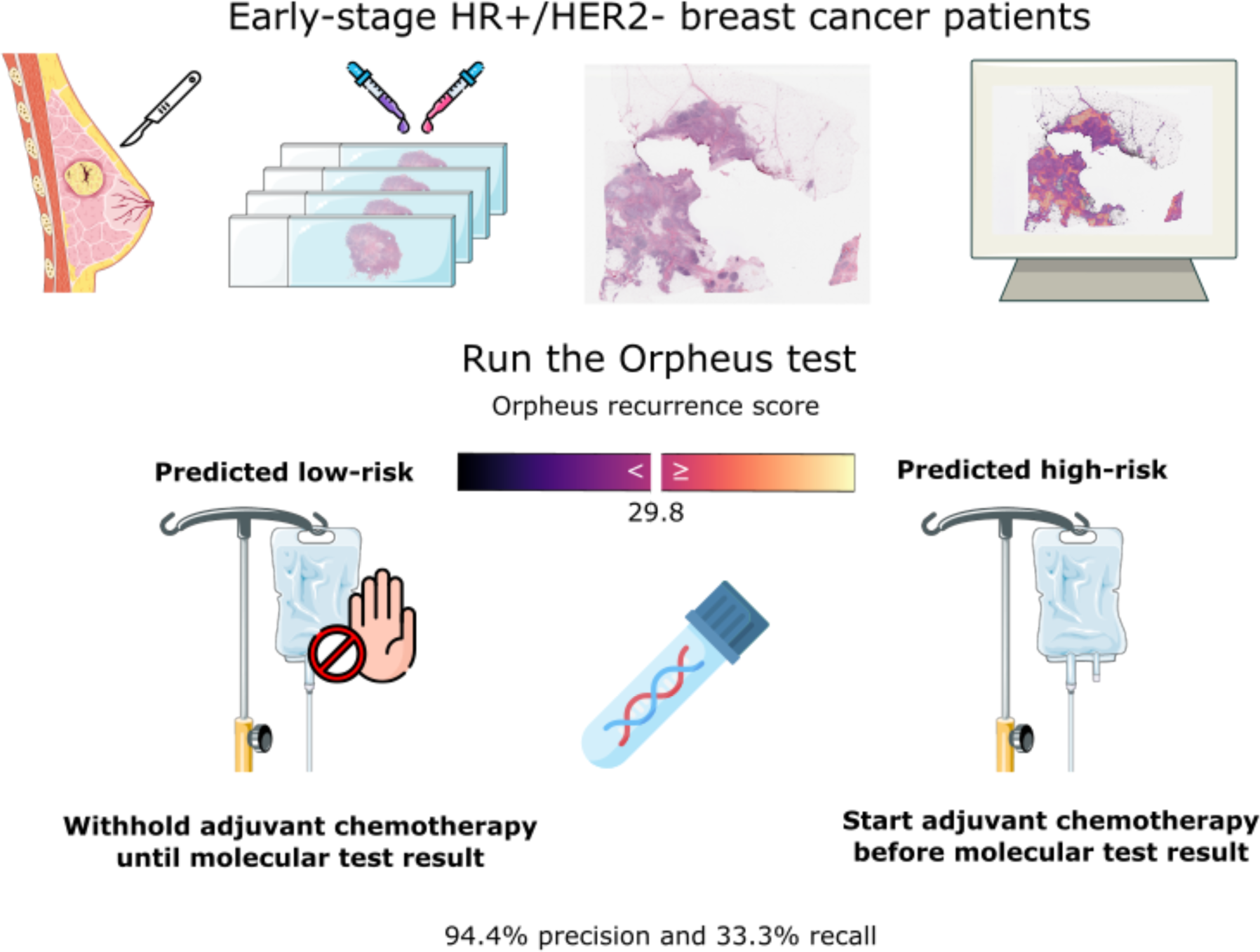
Potential clinical use-case of the Orpheus recurrence risk prediction model. The Orpheus multimodal prediction model for recurrence risk prediction is potentially capable of guiding decision-making for adjuvant cytotoxic chemotherapy alongside adjuvant endocrine therapy with 94.4% precision and 33.3% recall as measured on the withheld test set of the MSK cohort (n=2338). The model is within scope for early-stage hormone receptor positive (HR+) and HER2-breast cancer patients.

